## Supplemental Figures 1-5 for "Prime Editing Efficiency and Fidelity are Enhanced in the Absence of Mismatch Repair"

### Supplementary Information

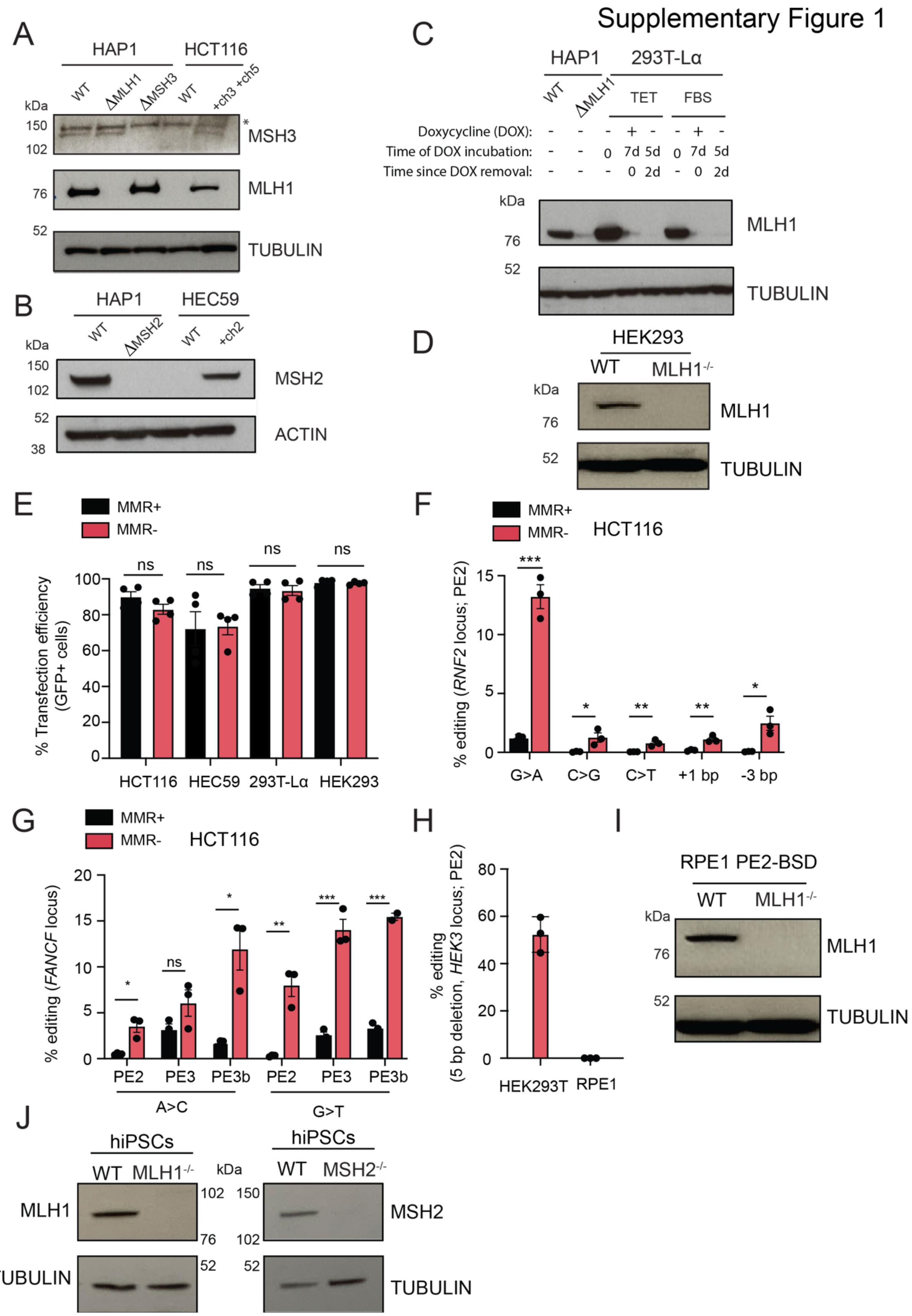

**Supplementary Figure 1: Characterisation of mismatch repair-proficient cell lines. A)** Immunoblot for MSH3, MLH1 and TUBULIN on cell extracts from HAP1 (WT, ΔMLH1, ΔMSH3) and HCT116 (WT and complemented with chromosome 3 and 5) cells. * denotes a non-specific band. **B)** Immunoblot for MSH2 and ACTIN in cell extracts from HAP1 (WT, ΔMSH2) and HEC59 (WT and complemented with chromosome 2). **C)** Immunoblot for MLH1 and tubulin in HAP1 (WT, ΔMLH1) and 293T-Lα cells. 293T-Lα were cultured in 10% of either Tet-approved FBS (TET) or regular foetal bovine serum (FBS). In both conditions, MLH1 is overexpressed without addition of doxycycline. Upon 7 days of doxycycline induction, MLH1 expression is abrogated. MLH1 abrogation persists 2 days after doxycycline removal. **D)** Immunoblot for MLH1 and tubulin in HEK293 WT cells, as well as an MLH1-isogenic knockout ($MLH1^{-/-}$). **E)** Transfection efficiency of the cell lines used, measured by flow-cytometry after transfection of a GFP-positive plasmid in at least three independent biological replicates. **F)** Efficiency of PE2 for different mutations in the *RNF2* locus, measured in HCT116 cells complemented with chromosomes 3 and 5 (MMR+), as well as HCT116 WT (MMR-). Values correspond to editing efficiency, measured by amplicon sequencing analysis in three independent biological replicates. **G)** Efficiency of P2, PE3 and PE3b after installation of an A>C or a G>T mutation in the *FANCF* locus, in HCT116 cells complemented with chromosomes 3 and 5 (MMR+), or HCT116 WT (MMR-). Values correspond to editing efficiency, measured by amplicon sequencing analysis in three independent biological replicates. **H)** PE2 efficiency after installation of a 5 bp deletion in the *HEK3* locus, in HEK293T cells as well as RPE1 cells. These cell lines express Cas9(H840A)-RT constitutively (PE2-BSD). Values correspond to editing efficiency, measured by Sanger sequencing and analysed by TIDE (Brinkman et al., 2014) in three biological replicates. **I)** Immunoblot of MLH1 and tubulin in RPE1 WT cells, as well an MLH1 isogenic knockout ($MLH1^{-/-}$).These cell lines express Cas8(H480)-RT constitutively (RPE1 PE2-BSD). **J)** Immunoblot for MLH1 (left) and MSH2 (right) in WT human induced pluripotent stem cells (hiPSCs) as well as isogenic knockouts for MLH1 (ΔMLH1) and MSH2 (ΔMSH2). Statistical analysis using unpaired t tests. Error bars reflect mean and SEM. Ns, p-value non-significant; *, p-value < 0.05.

Supplementary Figure 2

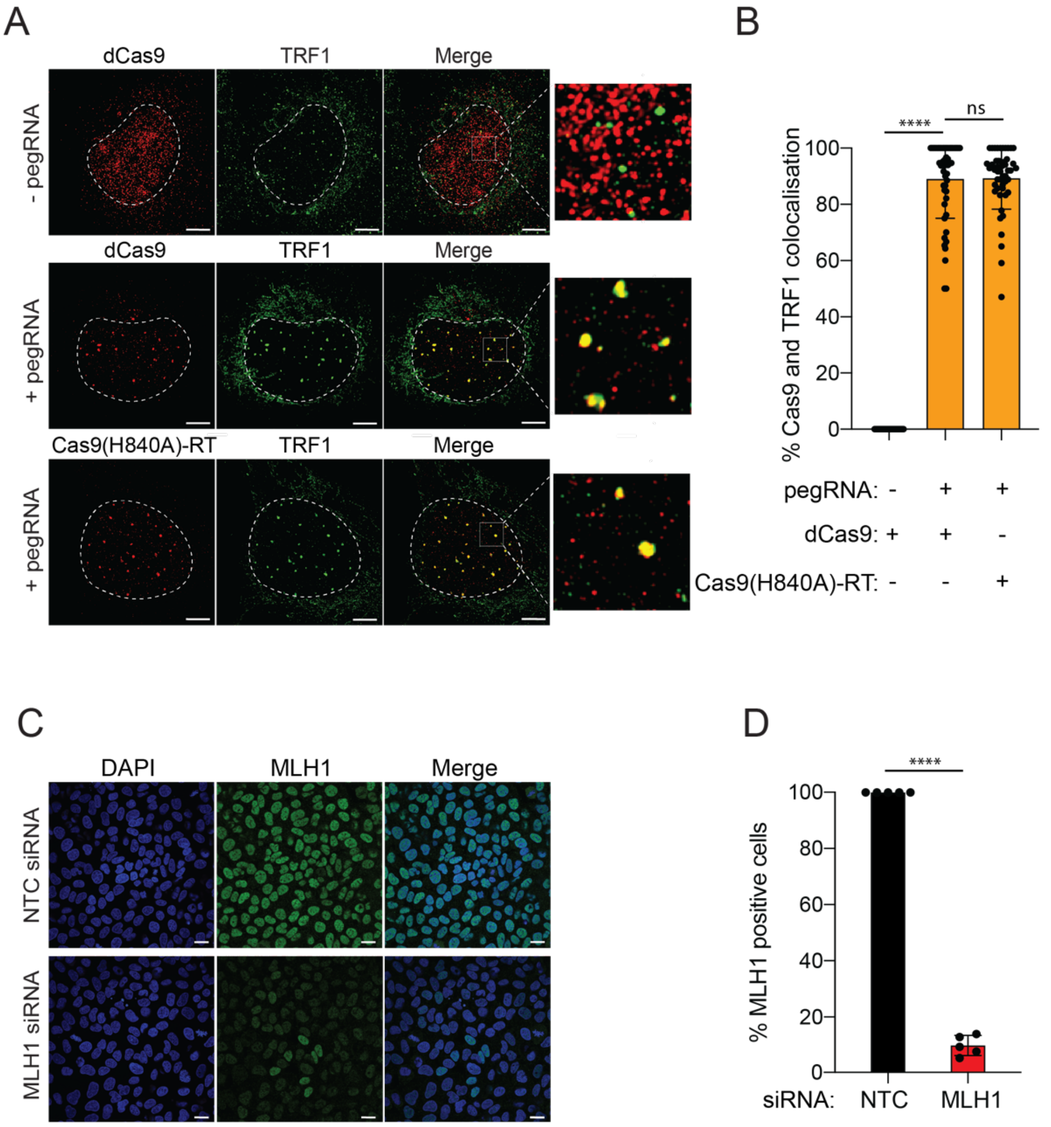


**Supplementary Figure 2: Localisation of proteins to sites of active prime editing. A)** Representative super-resolution images of dCas9, or Cas9(H840A)-RT, and TRF1 in U2OS cells, 24 hours following reverse transfection in the presence and absence of a pegRNA targeting telomeric repeats. Data from three biological replicates and at least 50 cells per condition. Scale bars = 5 µm. **B)** Quantification of A, indicating the percentage of dCas9 and Cas9(H840A)-RT foci that co-localise with TRF1, in the presence or absence of a pegRNA targeting telomeric regions. **C)** Representative images of MLH1 staining in U2OS cells transfected with either a non-targeting siRNA (NTC) or an siRNA targeting MLH1. Data from three biological replicates and over 500 cells per condition. Scale bars, 20 µm. **D)** Quantification of C indicating percentage of MLH1-positive cells. Statistical analysis using multiple unpaired t tests. Error bars reflect mean and SEM. Ns, p-value non-significant; ****, p-value < 0.0001.

#### Supplementary Figure 3

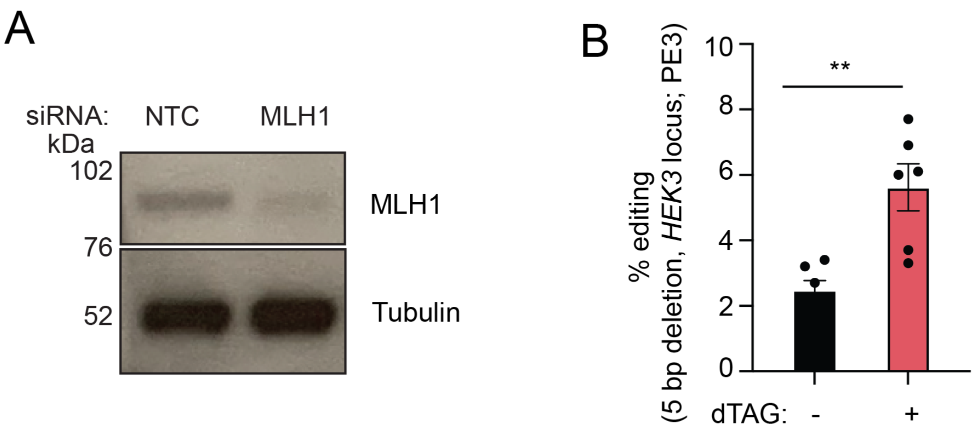


**Supplementary Figure 3: Strategies to improve prime editing efficiency by transient MLH1 ablation. A)** Immunoblot for MLH1 and tubulin in HEK293 cell extracts, three days post transfection with non-targeting control (NTC) or MLH1 siRNA pools (of four siRNAs). **B)** PE efficiency (PE3) of a 5 bp deletion in the *HEK3* locus in HAP1 dTAG-MLH1 cells in the presence and absence of 500 nM dTAG-7. Values correspond to editing efficiency, measured by Sanger sequencing and analysed by TIDE (Brinkman et al., 2014). Data from three independent biological replicates, with two technical replicates each. Statistical analysis with unpaired t tests. Error bars reflect mean and SEM **, p-value < 0.01.

Supplementary Figure 4

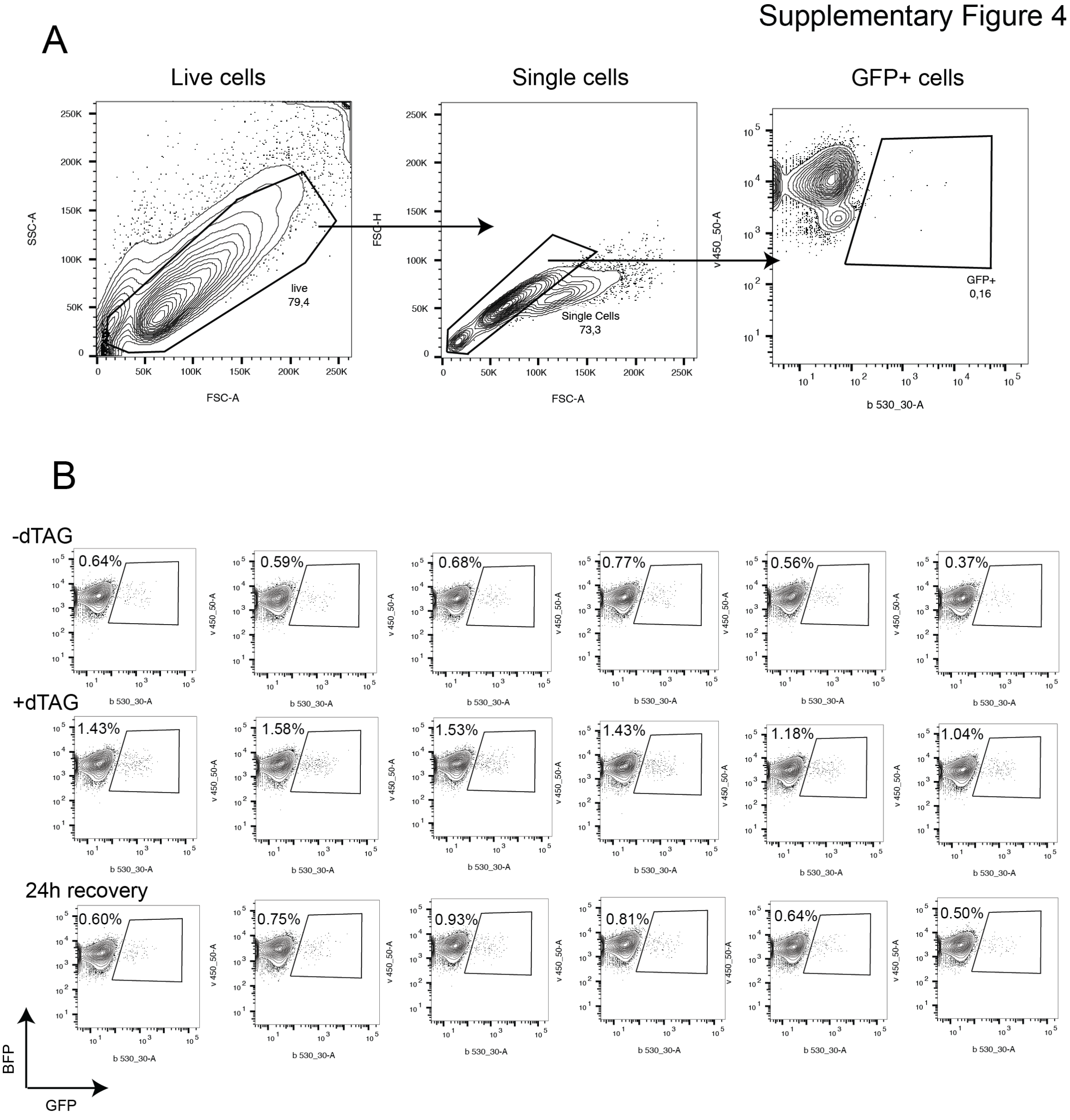


**Supplementary Figure 4: Gating strategy and flow-cytometry plots. A)** Gating strategy to assess BFP>GFP conversion by PE2. Live cells were gated on FSC-A and SSC-A profiles. Within this gate, doublets were excluded based on the FSC-A and FSC-H profile. BFP/GFP-double positive cells were gated within single cells, using a GFP-negative sample as a control. **B)** FACS plots showing the percentage of BFP>GFP conversion by PE in dTAG-MLH1 HAP1 cells, treated with the dTAG ligand ('+dTAG'), untreated ('-dTAG'), or after 24 hours in ligand-free media ('24h recovery'). The experiment was performed in three biological replicates, with two technical replicates each.

### Supplementary Figure 5

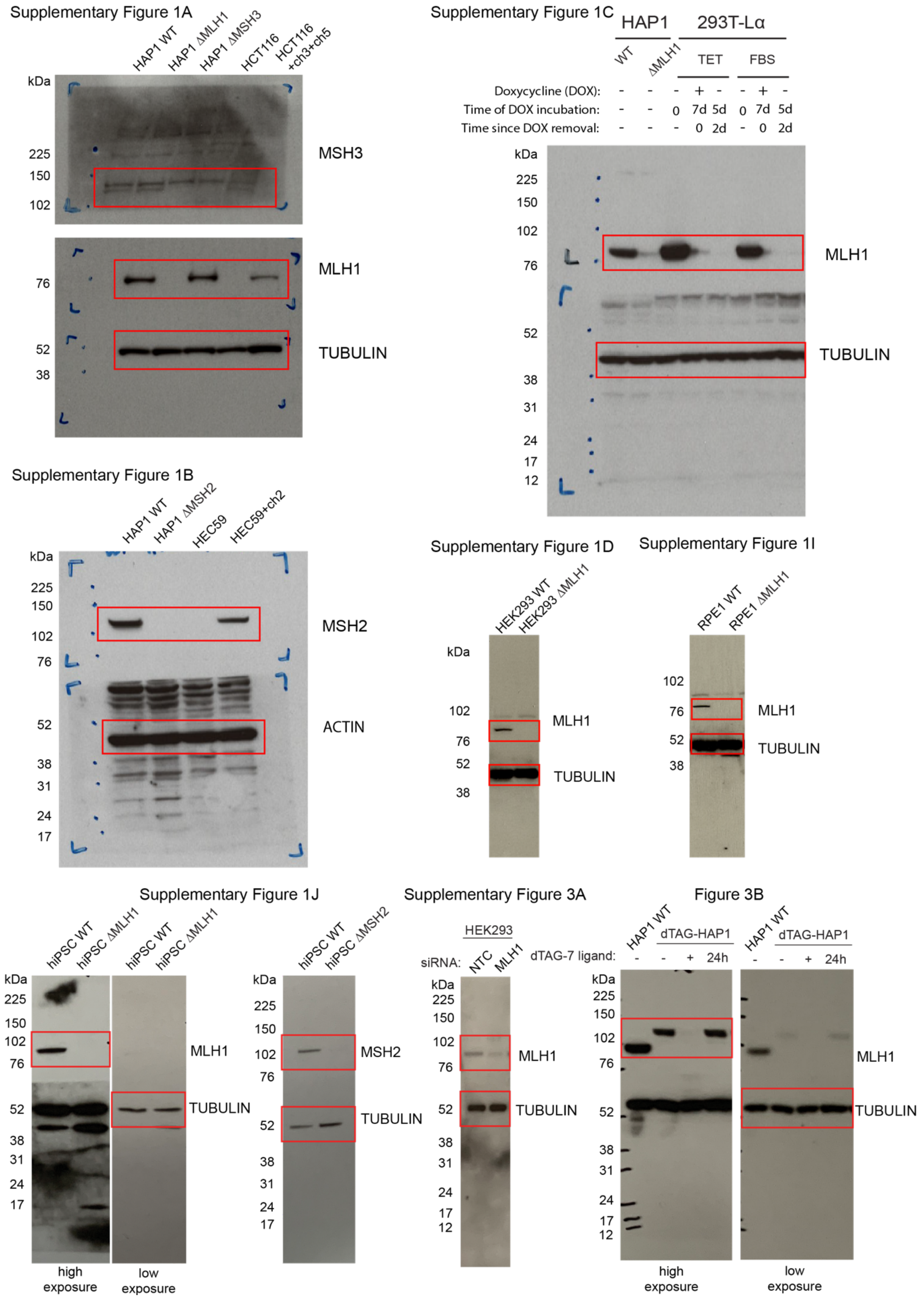


**Supplementary Figure 5: Uncropped immunoblots.**

**Description of Additional Supplementary Files**

File Name: Supplementary Data 1
Description: Frameshift mutations in collection of 32 knockout HAP1 cell lines, covering all DNA repair pathways.

File Name: Supplementary Data 2
Description: Sequences of pegRNAs, sgRNAs and primers used throughout the study
